# Integrative analysis of single-cell gene expression: A comprehensive database approach

**DOI:** 10.1101/2024.07.23.604709

**Authors:** Linh Truong, Thao Truong, Huy Nguyen

**Author notes:** **Correspondence** and requests for materials should be addressed to Huy Nguyen at. These authors contributed equally to this work.

## Abstract

The exponential growth of single-cell datasets provides unprecedented opportunities to advance our understanding of complex biological systems. However, effectively locating and integrating related studies for meaningful insights remains challenging. Traditional databases primarily index basic metadata, which necessitates time-consuming downloading and re-filtering based on gene expression and cell type or tissue composition, followed by computationally intensive aggregation. This process often results in excessively large datasets that are difficult to analyze effectively, further complicated by batch effects. To address these issues, we have developed a computational approach to efficiently extract and index both expression data and annotations. Our comprehensive database incorporates detailed author annotations and gene expression profiles, enabling refined searches and integrated analyses to uncover common biological patterns while accounting for the repeatability of patterns across multiple studies and mitigating batch effects. This approach significantly reduces computational demands and enhances the accessibility and utility of single-cell transcriptomics data for the broader research community. In the first version, we release a human database comprising 244 datasets from 236 cell types, 35 tissues, and 31 conditions.

## Introduction

The field of single-cell biology has undergone a revolutionary transformation, driven by the rapid development and widespread adoption of single-cell RNA sequencing (scRNA-Seq) technologies. These advancements have enabled researchers to dissect the complexities of biological systems at an unprecedented resolution, offering insights into the cellular heterogeneity that underlies physiological functions and disease states. However, the exponential growth in single-cell data generation introduces new challenges in accessing needed datasets and extracting human-understandable information. Traditional databases such as Gene Expression Omnibuse^1^ (GEO), the Single Cell Portal^2^, and CellxGene^3^ primarily focus on indexing limited metadata like study titles, abstracts, and categorical tags. Crucially, they lack the detailed annotation efforts from authors and the comprehensive expression profiles of cells, which are key to revealing biological mechanisms. Consequently, researchers often rely on keywords to navigate to studies relevant to their research. While newer databases like Single Cell Portal and CellxGene provide basic functionalities for examining gene expression and cell annotations within individual studies, researchers must still manually download each study for further integration to discern common patterns. This process not only demands substantial computational resources but also significant coding effort. Moreover, the resultant aggregate studies are often quite large, posing many challenges for later manual discovery. Without batch correction, the same population of cells is separated by batches, preventing the revelation of common patterns. Although batch correction might work well for populations with well-defined differences, closely related populations are often mixed together, preventing researchers from revealing the heterogeneity of the population, which plays a crucial role in understanding biological mechanisms. These challenges significantly slow the pace of discovery, despite the rapid accumulation of data.

In response to these challenges, we have developed a comprehensive database tailored for single-cell datasets. Significant efforts were made to standardize public datasets, harmonizing gene and cell annotations across different nomenclatures into a unified format. This standardization facilitates the alignment of disparate datasets, enabling more accurate comparative analyses. Alongside these efforts, we developed a computational approach to index massive datasets efficiently, allowing for rapid access to both author annotations and cell expression profiles. A key feature of our database is its user-friendly interface, designed to enable researchers to retrieve necessary information within seconds—a process that traditionally could take weeks of data processing. This accessibility is particularly beneficial for labs with limited computational resources, democratizing data access and analysis. Our database supports interactive engagement in three primary ways: (1) Study search: Researchers can immediately access a list of studies related to their research focus, utilizing comprehensive information from both author annotations and expression profiles. (2) Gene search: This function allows users to retrieve the expression patterns of target genes across different cell types and tissues. (3) Cell type search: Users can explore the characteristics of various cell types available in the database. The query results available through our database are drawn from hundreds of studies, enabling the identification of common biological patterns across vast datasets. Our first release focuses on human data and includes 244 publicly available datasets, comprising 236 cell types, 35 tissues, and 31 conditions. By providing a comprehensive and user-friendly database for single-cell datasets, we aim to accelerate the discovery rate and enhance our understanding of complex biological systems.

## Results

### Overview of the indexing workflow

The foundation of our database is built upon the meticulous work of countless scientists who have diligently studied, explored, and manually labeled their single-cell data. We aim to combine these values to reveal common patterns across multiple studies. While individual experiments may contain errors, recurring observations across independent studies from different labs indicate significant phenomena. These findings, even if not immediately explainable, can guide further experimental validation as hypotheses. In our approach, the number of studies that support a particular observation is more important than the number of cells within a single study that show the same result.

However, due to the independent nature of scientific research, variations in processing pipelines and reference alignments often lead to different gene annotations and cell population names. To address this challenge, we established a common control vocabulary to integrate the information consistently. For gene annotation, we used Ensembl gene annotations^4^ as a reference to convert various gene names, IDs, and aliases into a unified nomenclature. For cell annotation, we employed ontologies from EMBL-EMI^5^: Cell Ontology (CL) for cell type and subtype mapping, Human Disease Ontology (DOID) for condition mapping, and The BRENDA Tissue Ontology (BTO) for tissue mapping.

The workflow of our indexing pipeline is depicted in Figure 1, with a detailed description provided in the Methods section. In summary, we downloaded public datasets, including expression matrices and author annotations, and aligned them to our control vocabulary. After mapping, the expression of duplicated gene names is summed into one. The expression matrix was then processed to remove background noise, and gene expression was categorized as either expressed or not for each category in each study based on the author’s cell annotations. The indexed data were then merged into a single database, with each study contributing equally to the expressed/non-expressed status for gene-category relationships, regardless of the number of cells involved.

**Figure 1.**
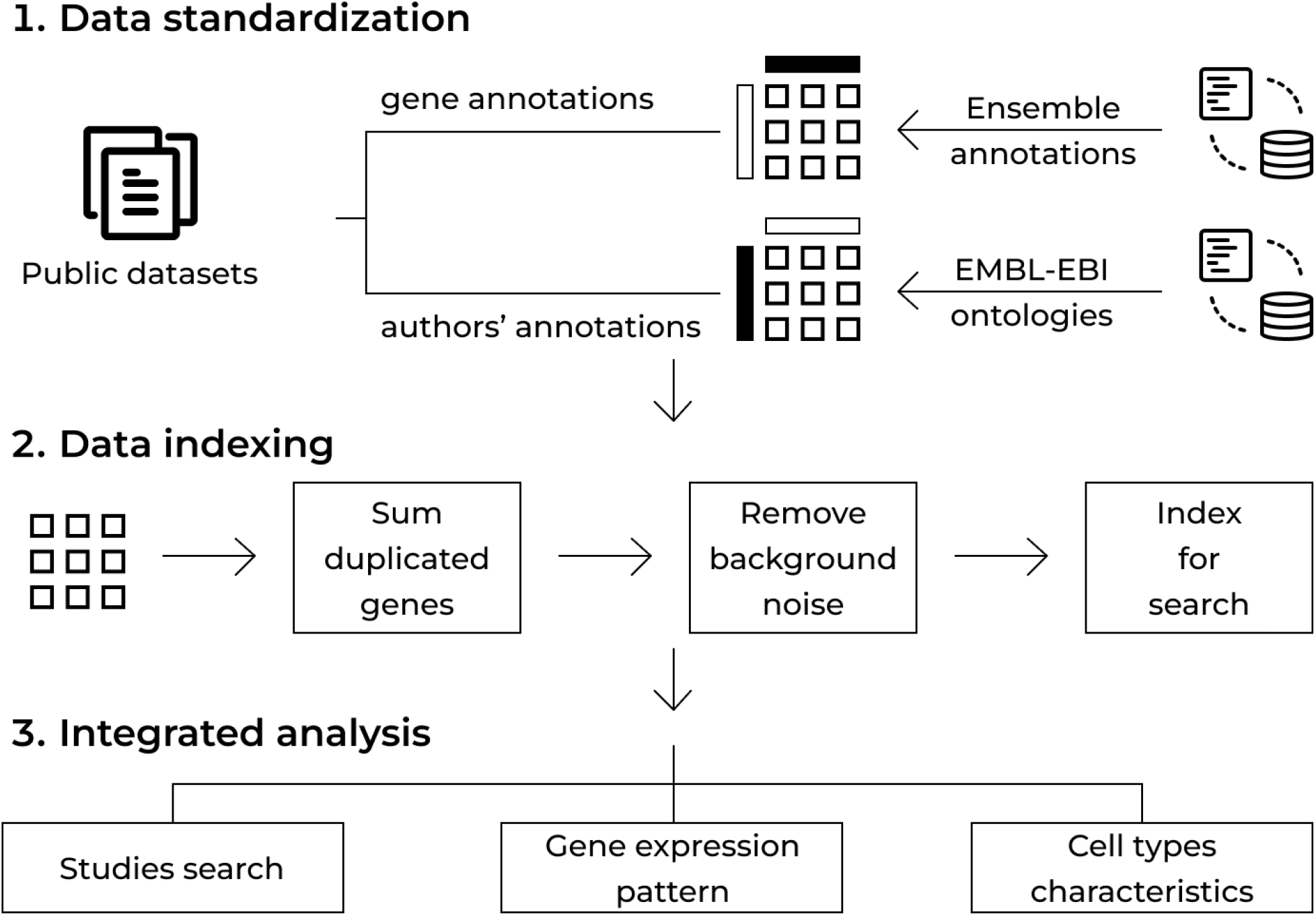
Schematic workflow of the indexing pipeline. The process of integrating and standardizing single-cell data involves several steps: (1) Data Standardization: Public datasets, including expression matrices and author annotations, are downloaded. Gene annotations are standardized using Ensembl gene annotations to unify gene names, IDs, and aliases. Cell annotations are standardized using EMBL-EBI ontologies, including the Cell Ontology (CL) for cell types, Human Disease Ontology (DOID) for conditions, and The BRENDA Tissue Ontology (BTO) for tissues. (2) Data Indexing: Duplicate gene expressions are combined, and the expression matrix is processed to eliminate background noise. Gene expression is then categorized as expressed or not for each category in each study based on the author’s annotations. The indexed data is merged into a single database. (3) Integrated analysis: Advanced computational analyses are performed to query necessary data and conduct comparative studies, such as identifying markers for specific cell types across the entire database.

Furthermore, due to the dropout characteristic of single-cell datasets, where zero values may not accurately represent the absence of expression but rather technological limitations, we also indexed the expression coverage for each gene across categories and datasets. This metric helps evaluate the impact of different technologies on gene detection in subsequent analyses. Finally, we applied computational approaches for querying necessary data and conducting comparative studies, such as identifying markers for specific cell types across the entire database.

### Studies search

To facilitate quick access to a list of studies meeting specific criteria, our database indexes all available information for each study, encompassing the following elements: (1) Study metadata: This includes the title, abstract, authors, all related deposited data access IDs, and data download sources, which are manually collected. (2) Cell metadata: All available annotations provided by the authors, and corresponding control vocabularies. (3) Gene expression: We determine if a gene is considered expressed in a study based on specific thresholds: the gene must be expressed in at least 100 cells and cover at least 15% of a cell population as per the author’s annotations (this can include any cell types, tissues, conditions, etc.). This threshold helps filter out random and lowly expressed events.

Studies can be queried using one of three methods or a combination of these methods: (1) Search by matched categories: This search method operates based on combinations of control vocabularies. Categories within the same ontology are queried using an OR operation, while categories across different ontologies are queried using an AND operation. For example, a query could be for studies of “lung tissues AND (T cells OR B cells).” (2) Search by gene expression:

This method allows extraction of a list of studies where a specific gene or list of genes is expressed. (3) Free text search: Researchers can input keywords for specific annotations from authors or novel cell populations that do not yet have official names. Keywords are matched against all available study metadata and author annotations (Figure 2A).

**Figure 2.**
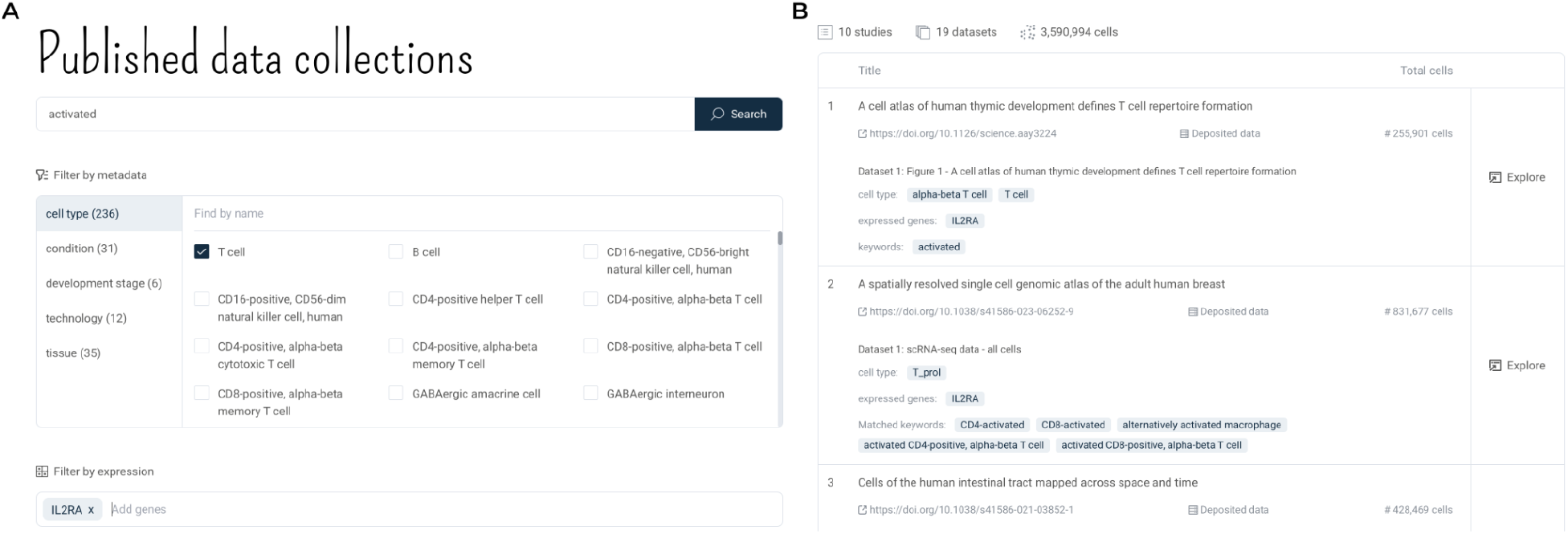
Studies search interface and results. (A) The search interface allows users to filter studies based on specific categories including cell types, conditions, development stages, technologies, and tissues. Users can perform a free text search or filter by gene expression, with selected filters displayed at the bottom of the panel. (B) The search results display a list of studies that match the specified criteria. Each entry includes the study title, publication DOI link, dataset description, cell types, expressed genes, matched keywords, and the total number of cells. The interface provides an “Explore” button for further investigation of each study’s details.

The list of matched studies is presented with the necessary information for further exploration, including study metadata, all available annotations from the authors, and the original annotations that matched the search queries (Figure 2B). This origin-matched information helps researchers quickly identify novel insights from other authors’ annotations.

### Genes search

Our database facilitates the exploration of gene expression patterns across cell types and tissues by indexing expression data for each labeled cell type and tissue within collected studies. We exclude data from cell populations smaller than 100 cells to minimize random noise. A gene is considered expressed in a particular label if it appears in at least 15% of that cell population. We track the repetition of gene expression across labels, allowing each study to contribute only one count if at least one of its composite datasets shows the gene expressed in that label. This method maintains data integrity and prevents overrepresentation. When researchers query a gene, we retrieve and visualize its expression across various cell types and tissues. Each cell population is represented by a dot, with the dot color indicating the number of supporting studies and the dot size reflecting the average expression coverage of the gene from those studies. Gene expression interpretations can be summarized as follows: (1) Specificity: The fewer cell types or tissues a gene is expressed in, the more specific it is considered. (2) Reliability: The greater the number of supporting studies, the more reliable the expression data. (3) Capture Rate: A higher expression coverage suggests that a gene is characteristic of certain cell types or tissues; conversely, low coverage may suggest the presence of a sub-population or limitations in detection technologies.

To improve the interpretability of gene expression patterns across cell types, we utilize the relationships defined by Cell Ontology and display expressed cell types as a sub-ontology. A complex sub-ontology indicates widespread gene expression across many cell types, while a simpler depiction suggests greater cell type specificity. This approach significantly enhances the readability and specificity of gene expression related to cell lineages (Figure 3A, Figure 3B). For tissues, due to the smaller number of categories, results are presented consecutively and sorted by the number of supporting studies. We also index information from bulk datasets, including GTEx and TCGA, allowing for comparison between single-cell datasets, normal bulk RNA datasets, and cancer bulk RNA datasets (Figure 3C, Figure 3D).

**Figure 3.**
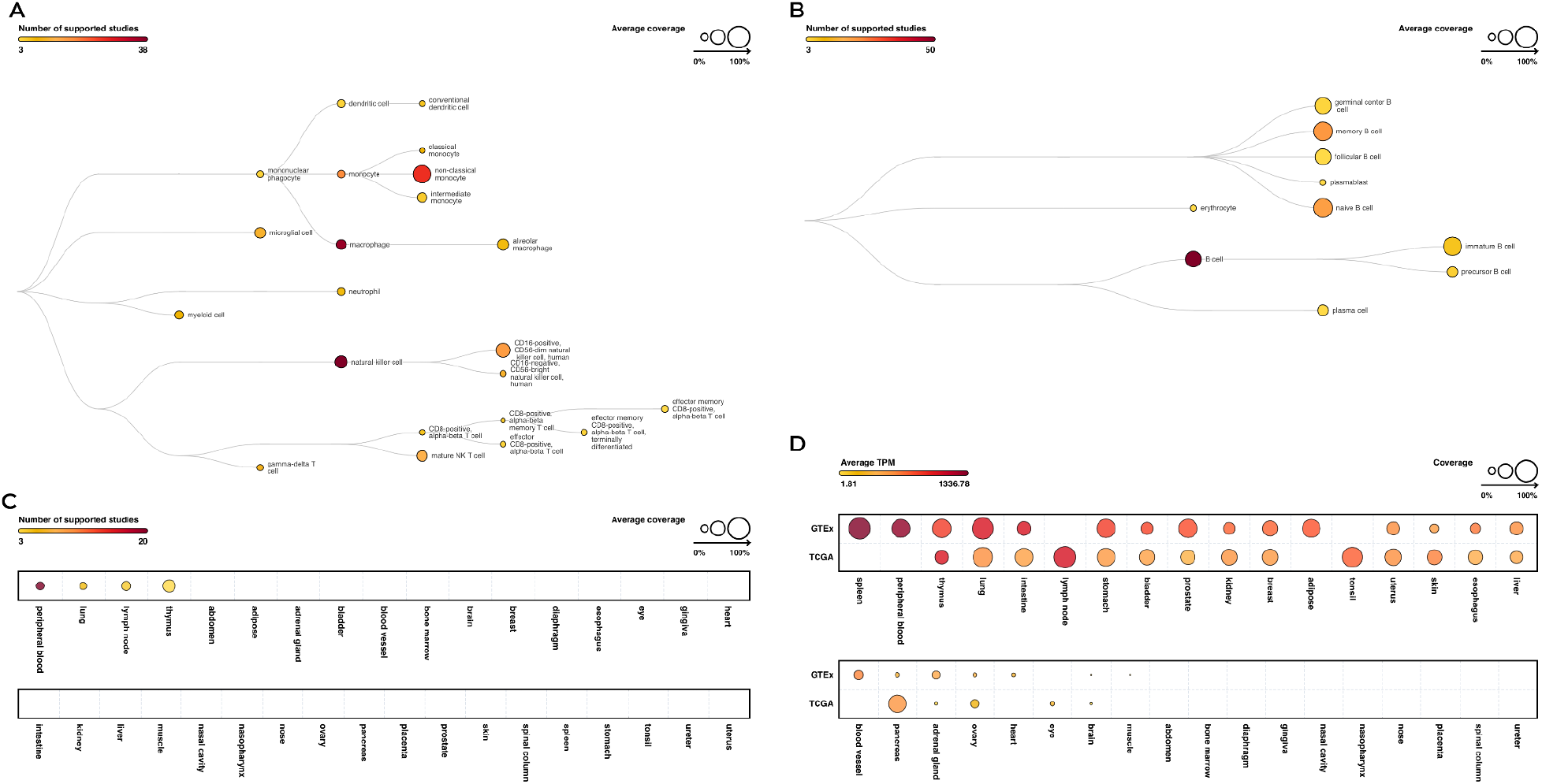
Visualization of gene expression patterns across cell types and tissues. (A) and (B) present the ontology-based visualization of gene expression patterns across various cell types. Each dot represents a cell population where the gene is expressed, with the dot color indicating the number of supporting studies and the dot size reflecting the average expression coverage from those studies. The complexity of the ontology corresponds to the gene’s expression specificity: a more complex ontology indicates widespread expression across many cell types, while a simpler depiction suggests greater specificity to particular cell types. (C) and (D) show gene expression patterns across tissues, sorted by the number of supporting studies. The results are enhanced by integrating data from bulk RNA datasets, such as GTEx and TCGA, allowing for comparisons between single-cell, normal bulk RNA, and cancer bulk RNA datasets.

In addition to expression data, we index detailed literature information about genes, including synonyms, protein products, descriptions, and related identifiers. This comprehensive indexing supports the understanding and explanation of expression patterns. Researchers can access all results in both tabular data and visual plots, with the option to retrieve a list of studies that support the findings directly from the data table.

### Cell types search

Our database facilitates the study of cell type characteristics by indexing specific markers for each cell type. Marker genes can be specific to a cell type and all its subtype lineages. Additionally, while cell subtypes within a lineage inherit these lineage markers, they also possess unique markers that differentiate them from other subtypes. Conversely, different cell lineages may share some markers, even as they are distinguished by major lineage-specific markers. Therefore, we systematically categorize all available cell types and subtypes from the indexed database into major groups based on lineage, using relationships defined by the Cell Ontology. Then, we employ a scoring system based on three criteria for evaluating a gene’s role as a specific marker: (1) Repetition: The number of studies that support the gene’s expression in that cell type—the higher the number, the stronger the evidence. (2) Major cell type specificity: The number of other major cell types that express the gene—the fewer the number, the more specific the marker. (3) Cell type specificity: The number of other cell types within the same major group that express the gene—the fewer the number, the more specific the marker. A gene is considered a marker of a cell type if it is supported by expression data from at least two studies. We rank genes according to a combined score from these criteria, and the top 30 results are retrieved for researchers studying cell type markers (Figure 4B). When researchers query a cell type, besides potential markers derived from analysis across the database, we also provide comprehensive literature information, including the name, synonyms, definition, and all available subtypes from our database. This information aids in further navigation to its subpopulations (Figure 4A). This data is indexed from the Cell Ontology.

**Figure 4.**
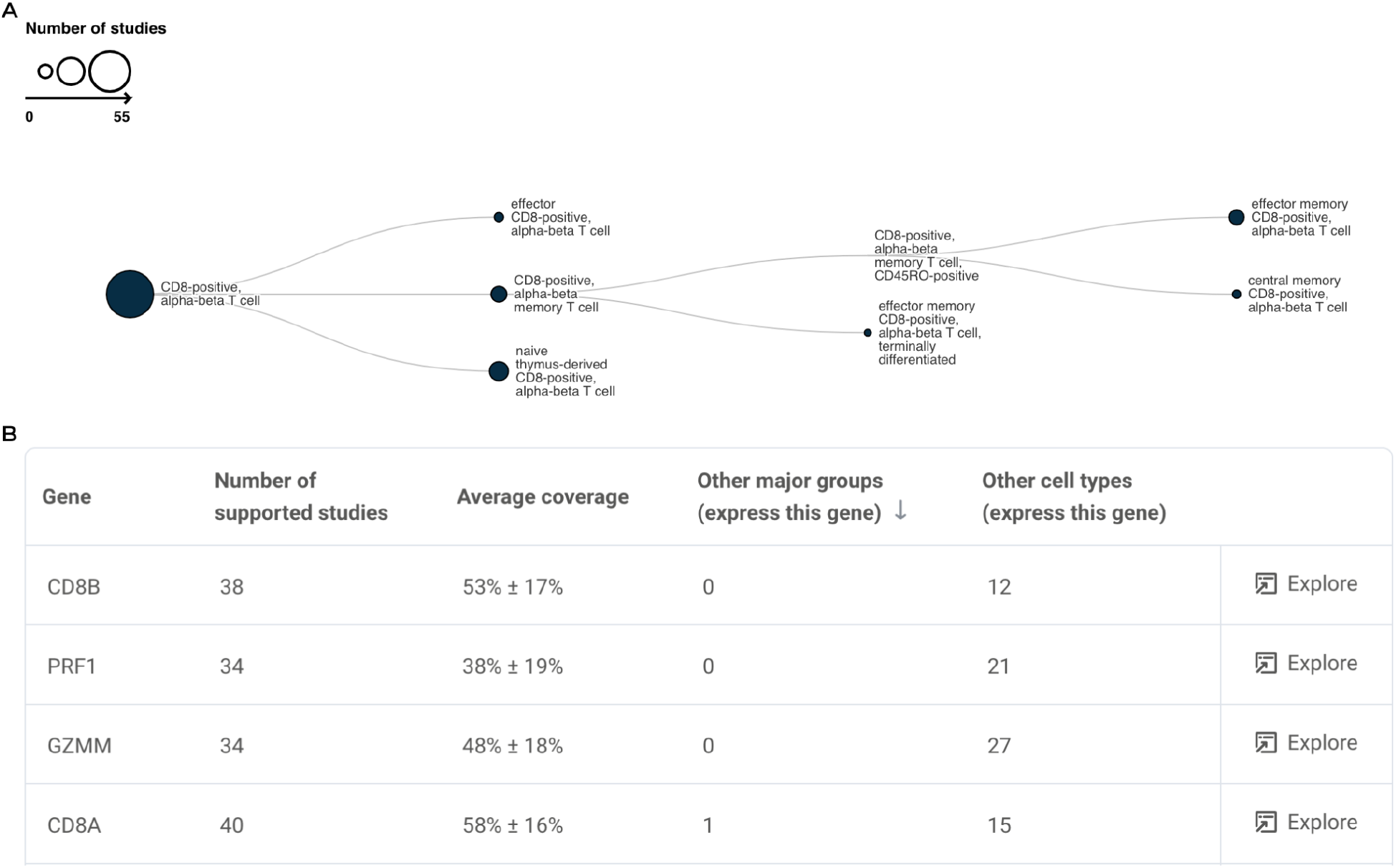
Cell type search results. (A) Visualization of cell type relationships using the Cell Ontology. Each node represents a cell type, with lines indicating lineage relationships. The size of each node reflects the number of studies supporting the expression of specific marker genes. (B) Tabulated results for marker gene search in a specific cell type. The table includes the gene name, the number of supporting studies, the average coverage percentage, the number of other major cell groups that express the gene, and the number of other cell types within the same major group that express the gene. The “Explore” button allows users to further investigate the supporting studies for each gene.

## Discussion

The development of our comprehensive single-cell database represents a significant advancement in the exploration and analysis of gene expression patterns across various cell types and tissues. By indexing extensive datasets and implementing user-friendly search functionalities, we enable researchers to efficiently access and interpret the wealth of information available in the field of single-cell biology.

One of the primary future goals for our database is the expansion of its scope. We plan to incorporate more studies and datasets, ensuring that our resource remains up-to-date and comprehensive. Furthermore, we aim to extend our database to include data from other species, allowing for comparative analyses and the identification of conserved and species-specific patterns. In addition to expanding the database’s content, we also plan to enhance its analytical capabilities. One such improvement involves the integration of higher-level analyses, such as co-expressed genes and comparative analyses between specific combinations of annotations. By identifying co-expressed genes, researchers can gain insights into potential gene interactions and regulatory networks, while comparative analyses will enable the discovery of shared and unique features across different cell types, tissues, and conditions.

In summary, our single-cell database serves as a valuable resource for researchers in the field, and we are committed to continuously improving and expanding its capabilities to meet the evolving needs of the scientific community.

## Methods

### Raw data collection and processing

Public single-cell datasets were obtained from public repositories, including GEO, Single Cell Portal, and CellxGene. When datasets were available in multiple repositories, CellxGene was preferred as it hosts the fully processed data with annotations as Scanpy/Seurat objects. We developed different pipelines to convert various data formats (tsv, csv, mtx, hdf5, scanpy, Seurat) into a unified data structure for the indexing pipeline.

During data format conversion, we mapped different gene IDs and gene aliases to controlled gene names using annotations from Ensembl release version 111 (https://ftp.ensembl.org/pub/release-111/). For human gene annotations, we combined information from GRCh37 and GRCh38 references, mapping old gene names from GRCh37 to current gene names from GRCh38 using Ensembl IDs. Gene alias information was also retrieved from the UniProt database to map name aliases to a common name from Ensembl. Expression values for genes with the same name were summed into a single row.

Background noise was removed for each gene by calculating the 90th quantile of expression across all cells. Expression values that were 10 times less than this threshold were removed.

### Curation process for cell annotation

For each study, we extracted all author annotations and manually navigated through each study to select a proper field from the author annotations to map to the control vocabularies of cell types, tissues, conditions, developmental stages, and technologies.

Control vocabularies for cell types, tissues, and conditions were retrieved from the EMBL-EBI website. Cell Ontology (CL) was used for cell type and subtype mapping, Human Disease Ontology (DOID) for condition mapping, and The BRENDA Tissue Ontology (BTO) for tissue mapping.

After finishing standardizing all studies, we further categorized all available cell types and subtypes into major groups based on the relationship from Cell Ontology.

### Indexing process

Each gene’s expression was indexed as either expressed or not for each cell type and tissue label within the studies. A gene was considered expressed in a cell if its expression was above the background noise threshold. A cell population was considered to express a gene if at least 15% of its cells showed expression. A study was noted as expressing a gene if at least one cell type or population within it showed expression. Throughout this process, the expression coverage of each gene for each population and study was also recorded. Literature information about genes was retrieved from the UniProt database and indexed in JSON format Study information is indexed for searching, including title, abstract, authors, all related deposited data access IDs, and data download sources. This information is manually collected from the corresponding papers. We also design a data structure to efficiently index all author annotations and gene expression relationships to studies.

Cell-type literature information and relationships are retrieved from Cell Ontology and indexed in JSON format. Cell-type markers are pre-calculated and indexed for later queries. The calculation process is: (1) Extract all expressed genes per cell type/cell subtype, with a gene considered expressed in a cell type if more than 3 studies support the expression of that gene in that cell type. (2) Score the gene list: each gene in each cell type is scored using 3 criteria with descending weight: number of studies supporting the expression, number of other major cell types expressing this gene, number of other cell types in the same major group with this cell type expressing this gene. (3) Sort the gene list by the score and extract the top 30 results as potential markers. (4) Index the result for queries.

### Hosting Server Implementation

We designed and implemented a backend server for serving queries and a user-friendly website for easy interaction and information retrieval. The interface is designed as a set of actions, allowing researchers to retrieve all related information needed for exploration and visualization with just a few simple clicks.

## Supporting information

Supplementary Data 1: Studies search tutorial

Supplementary Data 2: Genes search tutorial

Supplementary Data 3: Cell types search tutorial

## Data availability

All data sets used in this work are available from publicly available sources. User-friendly web tools are freely available at https://omnibusx.com/.

## Acknowledgments

We thank OmnibusX Company Limited for supporting this research and providing access to the resources necessary to complete the project.

## Author contributions

Conceptualization, methodology, data curation and analysis, manuscript writing and coding: Linh Truong, Thao Truong, and Huy Nguyen.

## Competing interests

The authors declare no competing interests.

## Additional information

### Supplementary information

Supplementary Information is available for this paper.

## Notes

### Competing Interest Statement

The authors have declared no competing interest.

### Summary of Updates

The author's stage name has been replaced with their country original name

