## Supplementary Data 1: Studies search tutorial for "Integrative analysis of single-cell gene expression: A comprehensive database approach"

omnibusx
FROM DATA TO DISCOVERY

### Studies search

User guide for OmnibusX web tools

### Introduction

The field of single-cell biology has undergone a revolutionary transformation, driven by the rapid development and widespread adoption of single-cell RNA sequencing (scRNA-Seq) technologies. However, the exponential growth in single-cell data generation introduces new challenges in accessing needed datasets and extracting human-understandable information. Traditional databases focus on indexing limited metadata like study titles, abstracts, and categorical tags. Crucially, they lack the detailed annotation efforts from authors and the comprehensive expression profiles of cells, which are key to revealing biological mechanisms. Consequently, researchers often rely on keywords to navigate to studies relevant to their research, then manually download each study for further integration to discern common patterns. This process not only demands substantial computational resources but also significant coding effort.

Addressing this need, our database offers an advanced search functionality that encompasses not just traditional study information but extends to detailed author annotations and comprehensive expression profiles. This capability sets our platform apart from conventional databases, which typically limit searches to basic metadata like titles and abstracts. With our enhanced search function, researchers can efficiently sift through extensive datasets to find studies that are directly relevant to their specific scientific queries.

This guide will provide you with detailed instructions on how to utilize the Studies Search to its full potential, facilitating a smoother and more effective research process.

### Studies search

You can access the search tool directly at [https://omnibusx.com/studies](https://omnibusx.com/studies).

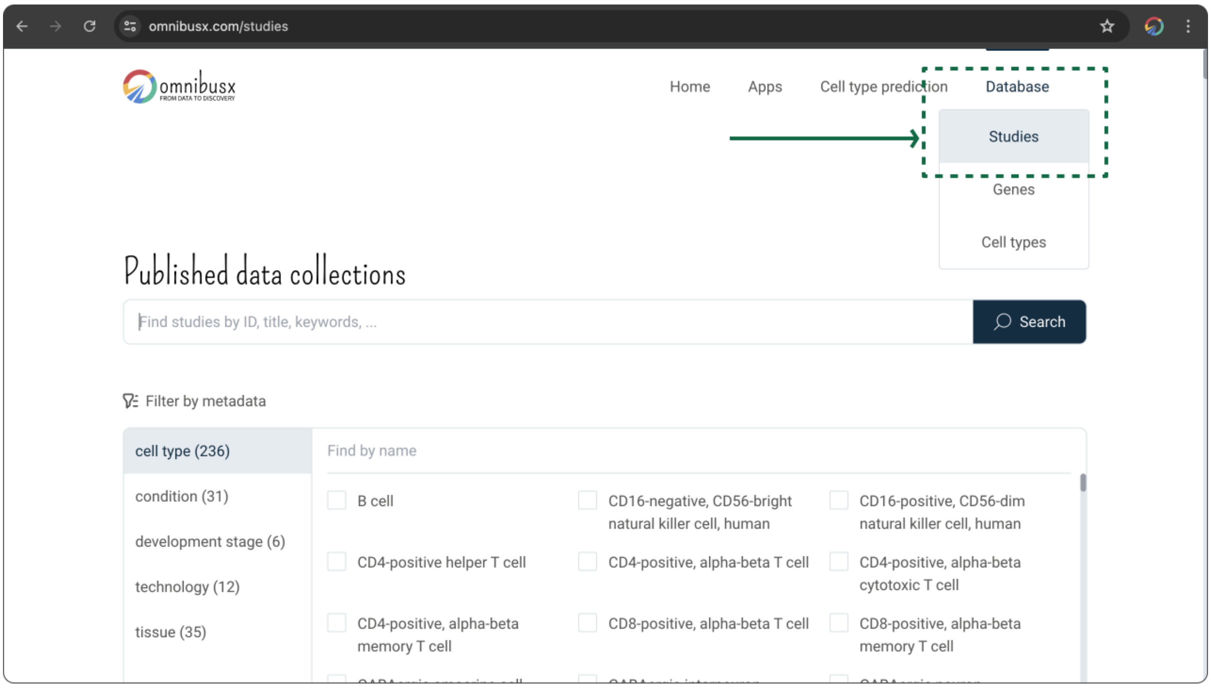


#### 1. Queries

##### 1.1. Free text search

To perform a free text search, enter your keyword in the text input and press `Enter` or click on the `Search` button. Your search will encompass all available text information related to a study, including titles, abstracts, author names, repository access IDs, and particularly, all annotations provided by the authors.

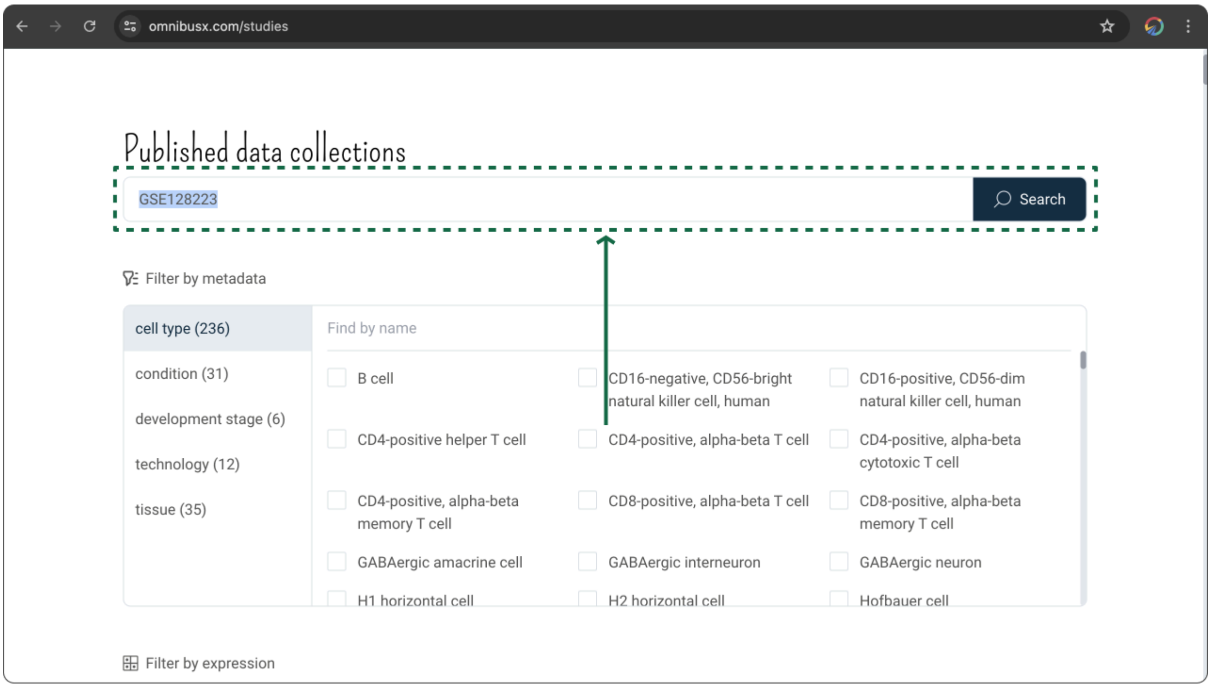


This function is especially useful for quickly locating a study by using well-known information such as the title or ID, or discovering novel annotations related to emerging cell populations that may not yet have official classifications.

#### 1.2. Controlled vocabularies search

Free text search might lead to missed results due to variations in naming conventions (e.g., CD4 T cell, CD4-positive T cell, CD4+). To accommodate naming variances, OmnibusX has extensively mapped different terminologies to standardized controlled vocabularies. You can utilize this feature by clicking on the needed categories from the `Filter by metadata` panel.

Selecting multiple categories from the same group (e.g., cell types like B cell or T cell) will apply an `OR` search filter, returning studies containing any of the selected categories. Selecting categories from different groups (e.g., tissue type and cell type) will apply an `AND` search filter, returning studies that meet all selected criteria (e.g., studies from lung tissues that contain B cells or T cells).

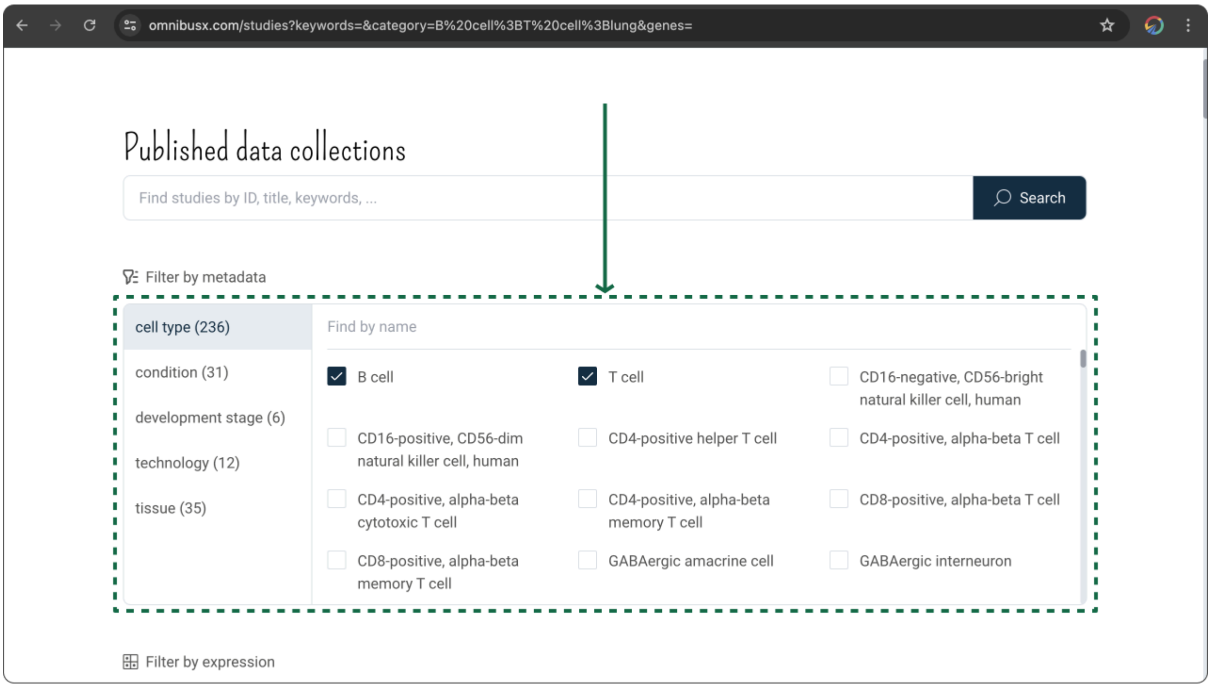


You can quickly find and select your target categories by typing the name into the filter input box above the metadata panel.

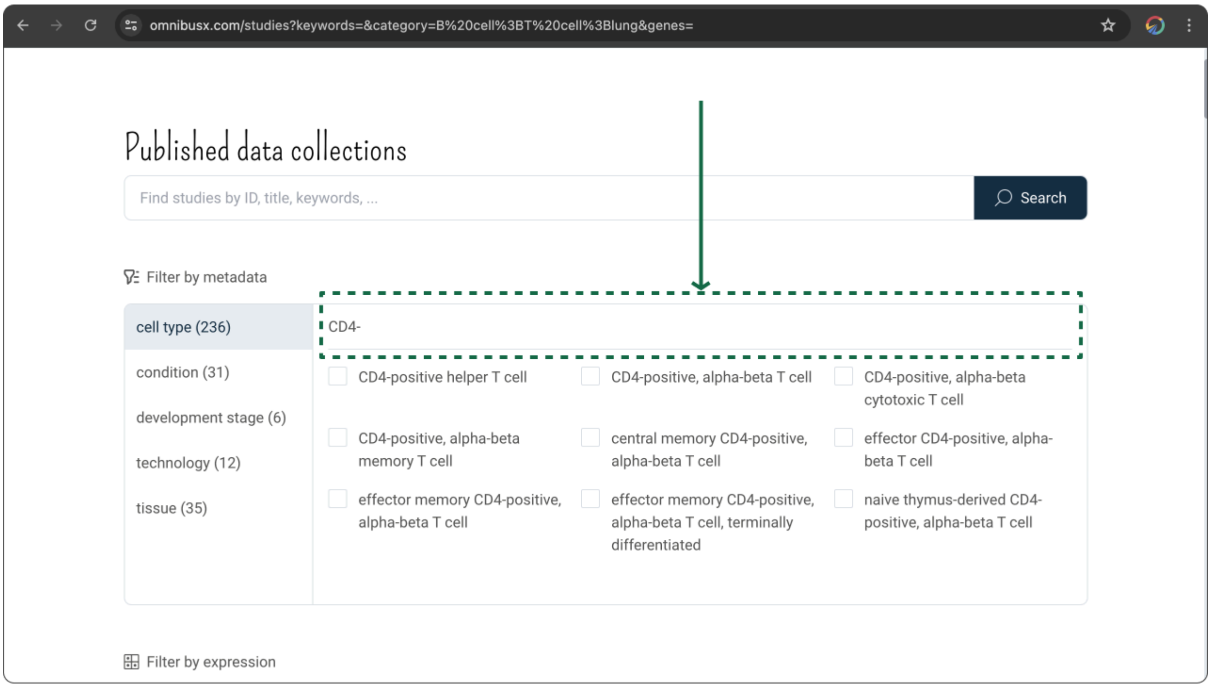

#### 1.3. Expression search

This advanced feature of OmnibusX allows you to efficiently locate studies that report the expression of specific genes, saving significant time that would otherwise be spent downloading and processing datasets to verify gene expression. To use this feature, input the gene name into the search box in the `Filter by expression` section. As you type, OmnibusX will suggest related genes which you can select directly.

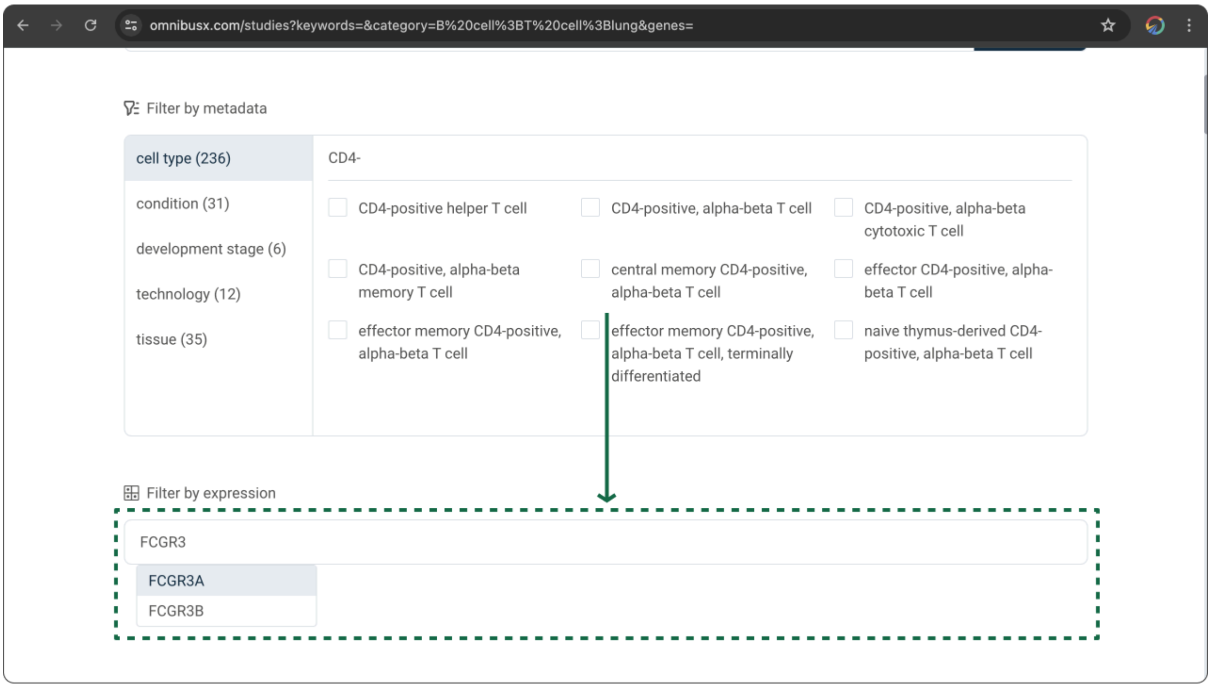


### 2. Results

Once your search is executed, all matching studies will be displayed in a tabular format. The number above the table provides a summary of the total search results. The table enables quick access to basic information for each study, including the title, reference URL, data deposited URL, and the number of cells. For each dataset associated with a study, all queries matching your search criteria will be listed for easy review. Original author annotations are displayed rather than controlled vocabularies to highlight potentially novel annotations that may be of interest. You can click the `Explore` button associated with each study to delve deeper into detailed data and further investigate the study's specifics.

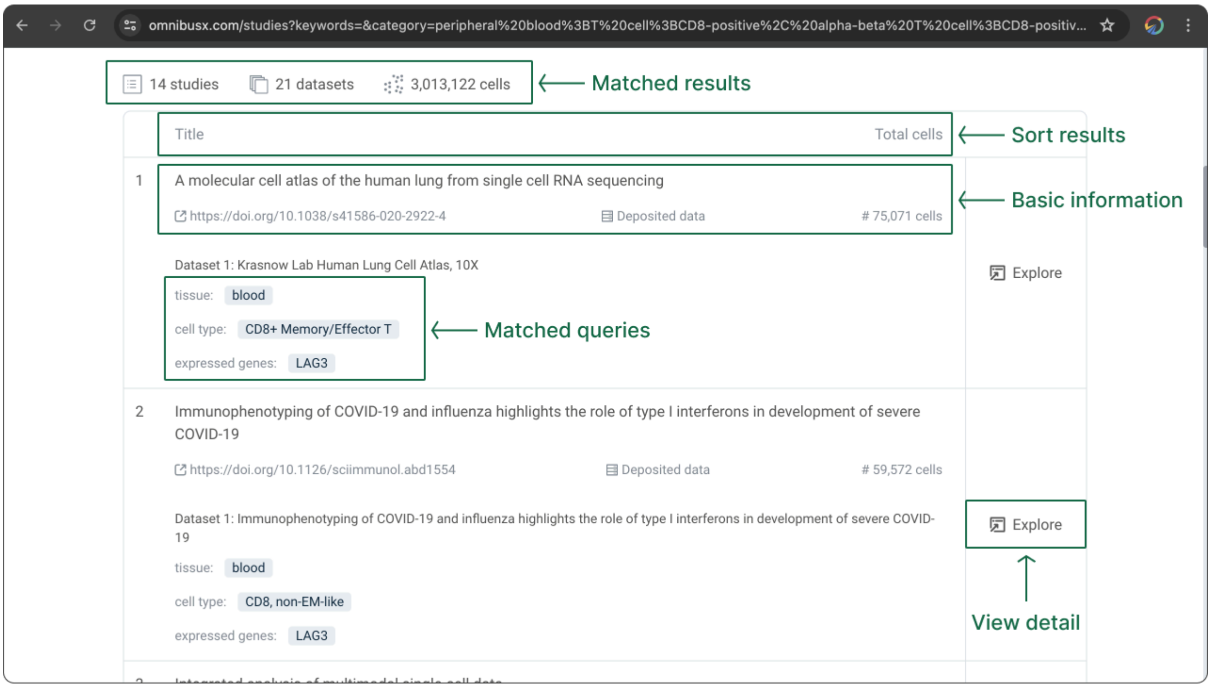


#### 3. Explore

On selecting the `Explore` option for a specific study, you will be presented with comprehensive information about that study. This includes:

- `Title`, `Authors`, and `Abstract`: Fundamental details providing a quick overview of the study's scope and purpose.

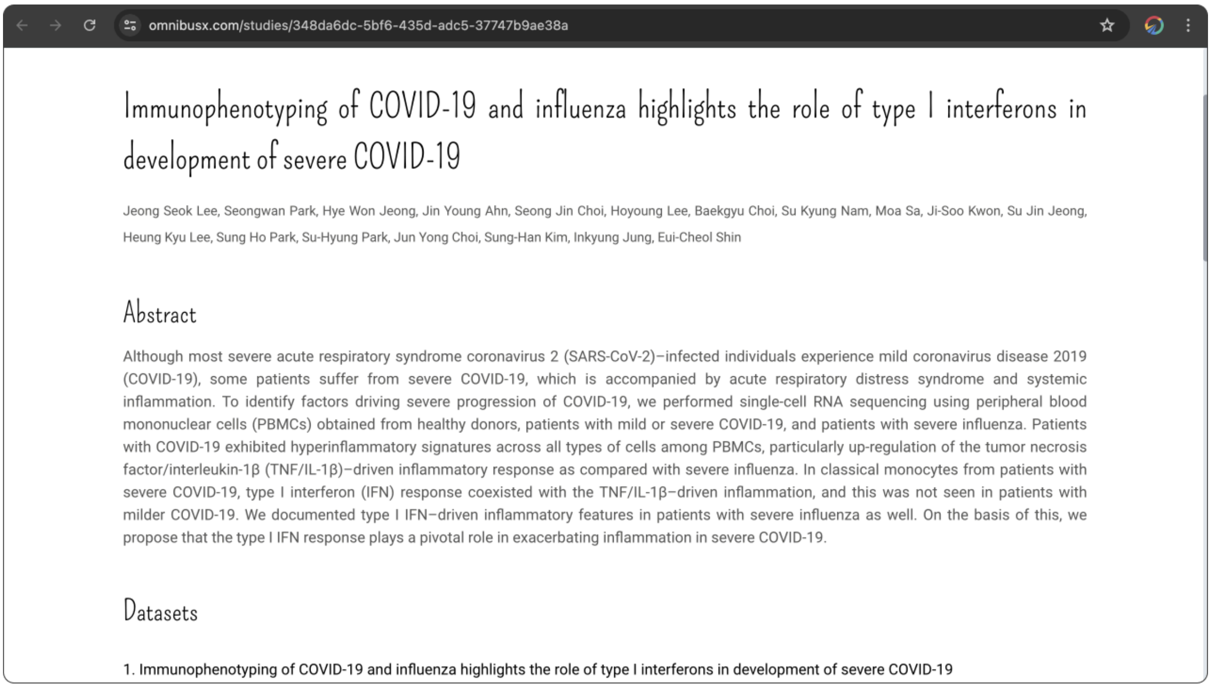


- `Annotations`: All available annotations from the authors are accessible. This section allows you to toggle between different fields from the summary table to explore various aspects of the study in greater depth.

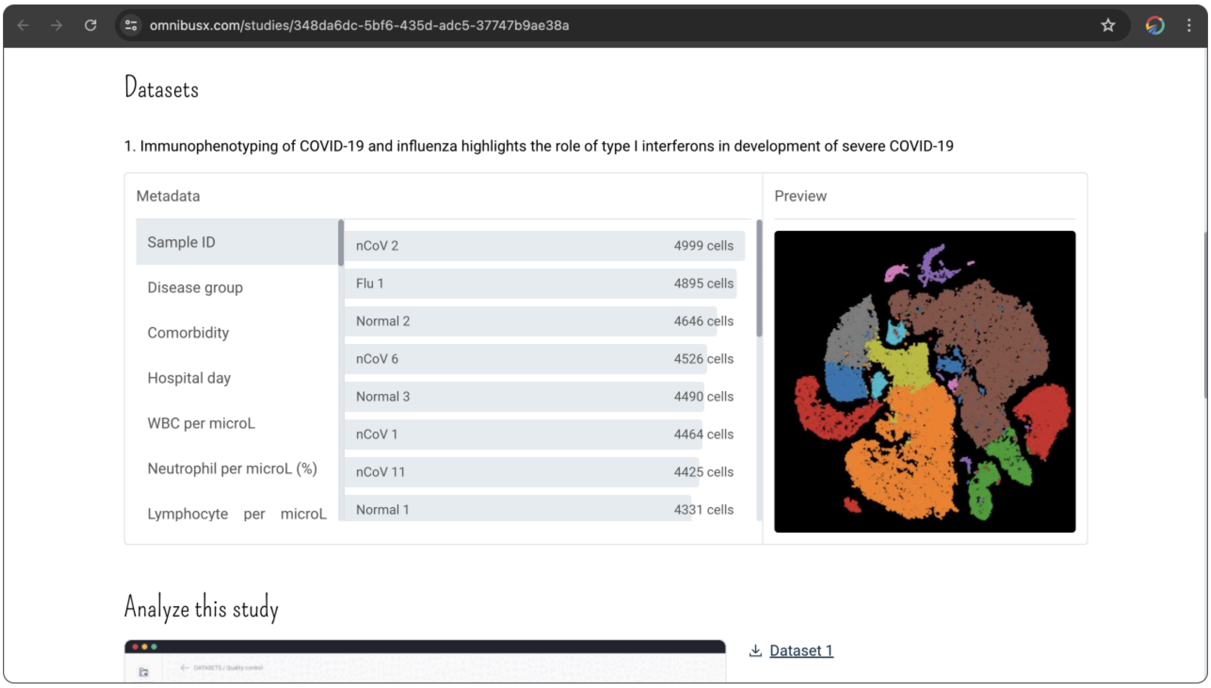


- `Data source` and `Accession IDs`: Direct links to the original data sources and relevant accession IDs, facilitate easy access to raw data.

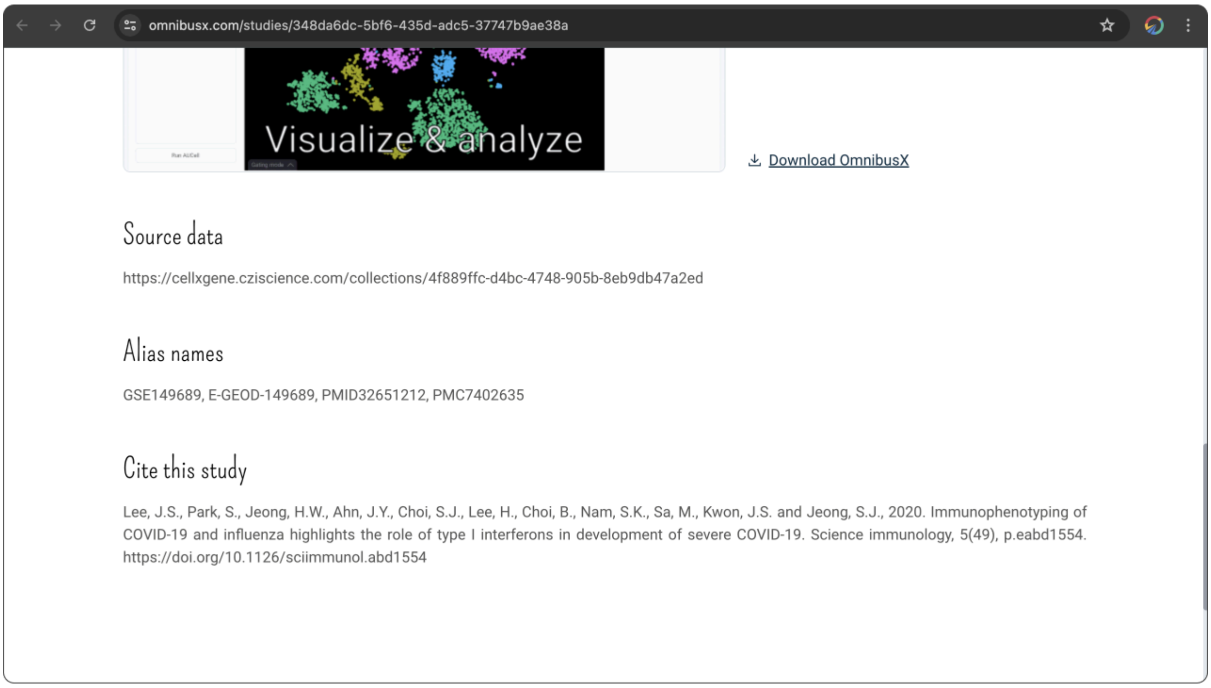
omnibusx.com/studies/348da6dc-5bf6-435d-adc5-37747b9ae38a
Visualize & analyze
Download OmnibusX
Source data
https://cellxgene.cziscience.com/collections/4f889ffc-d4bc-4748-905b-8eb9db47a2ed
Alias names
GSE149689, E-GEOD-149689, PMID32651212, PMC7402635
Cite this study
Lee, J.S., Park, S., Jeong, H.W., Ahn, J.Y., Choi, S.J., Lee, H., Choi, B., Nam, S.K., Sa, M., Kwon, J.S. and Jeong, S.J., 2020. Immunophenotyping of COVID-19 and influenza highlights the role of type I interferons in development of severe COVID-19. Science immunology, 5(49), p.eabd1554. https://doi.org/10.1126/sciimmunol.abd1554

### Thank you!

We extend our heartfelt gratitude to all users of the OmnibusX Studies Search platform. Your engagement and feedback are invaluable to us and are what drive continuous improvement and innovation within our database. We are committed to supporting the scientific community by providing robust tools that facilitate groundbreaking research and discovery.

If you have suggestions, feedback, or would like to share how OmnibusX has assisted in your research endeavors, please do not hesitate to reach out to us at. Your stories inspire us, and your feedback helps us refine our tools to better serve your needs.
