## Supplementary Data 2: Genes search tutorial for "Integrative analysis of single-cell gene expression: A comprehensive database approach"

omnibusx
FROM DATA TO DISCOVERY

### Genes search

User guide for OmnibusX web tools

### Introduction

Single-cell RNA sequencing (scRNA-Seq) technologies have revolutionized the field of single-cell biology, providing unprecedented insights into the complexities of biological systems. However, the rapid growth of data generation has presented significant challenges in data management and utilization. Researchers often face difficulties in accessing and integrating vast datasets, requiring extensive computational resources and coding effort.

To address these challenges, we have developed a comprehensive database with advanced computational tools that efficiently index massive datasets, enabling swift access to author annotations and detailed cell expression profiles. Our user-friendly interface allows researchers to quickly retrieve the expression patterns of target genes across various cell types and tissues, significantly reducing the time required for data processing.

The Gene Search tool, powered by our extensive database, helps identify common gene expression patterns across cell types and tissues. This guide will help you fully leverage the capabilities of the Gene Search tool to enhance your research outcomes.

### Genes search

You can access the search tool directly at https://omnibusx.com/genes.

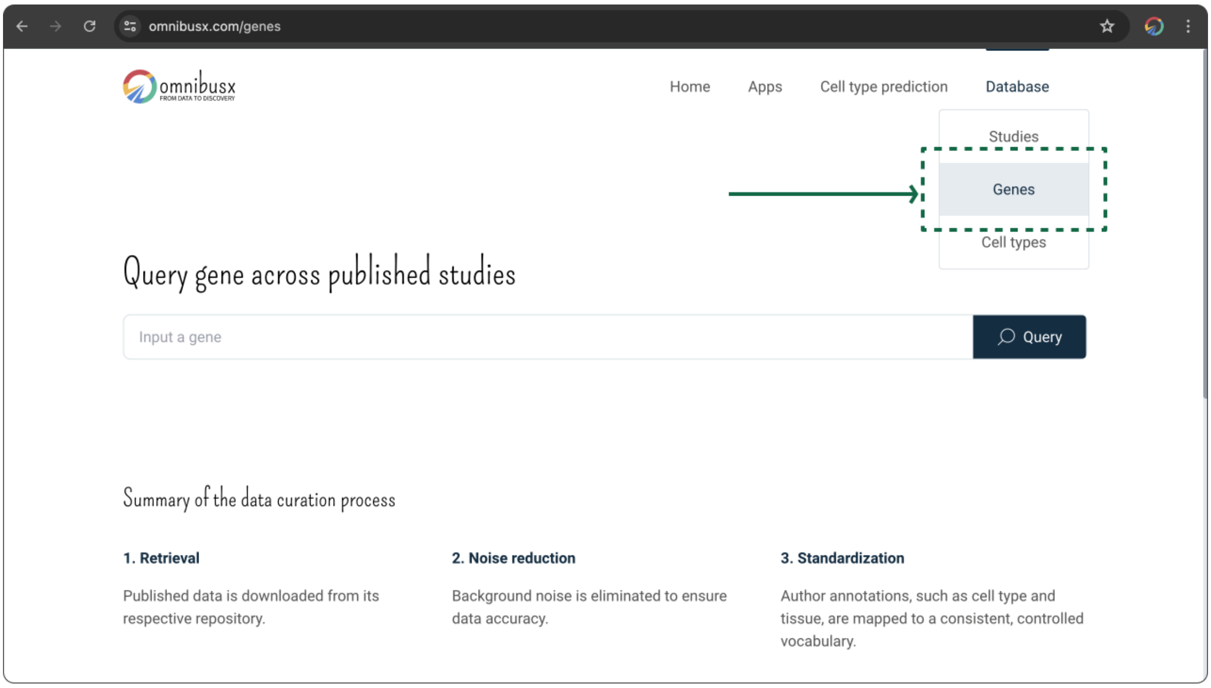


#### 1. Queries

You can begin by entering the target gene name into the input box on the `Gene search` page. As you type, OmnibusX will dynamically suggest related gene names, allowing you to select the correct one directly from the dropdown menu.

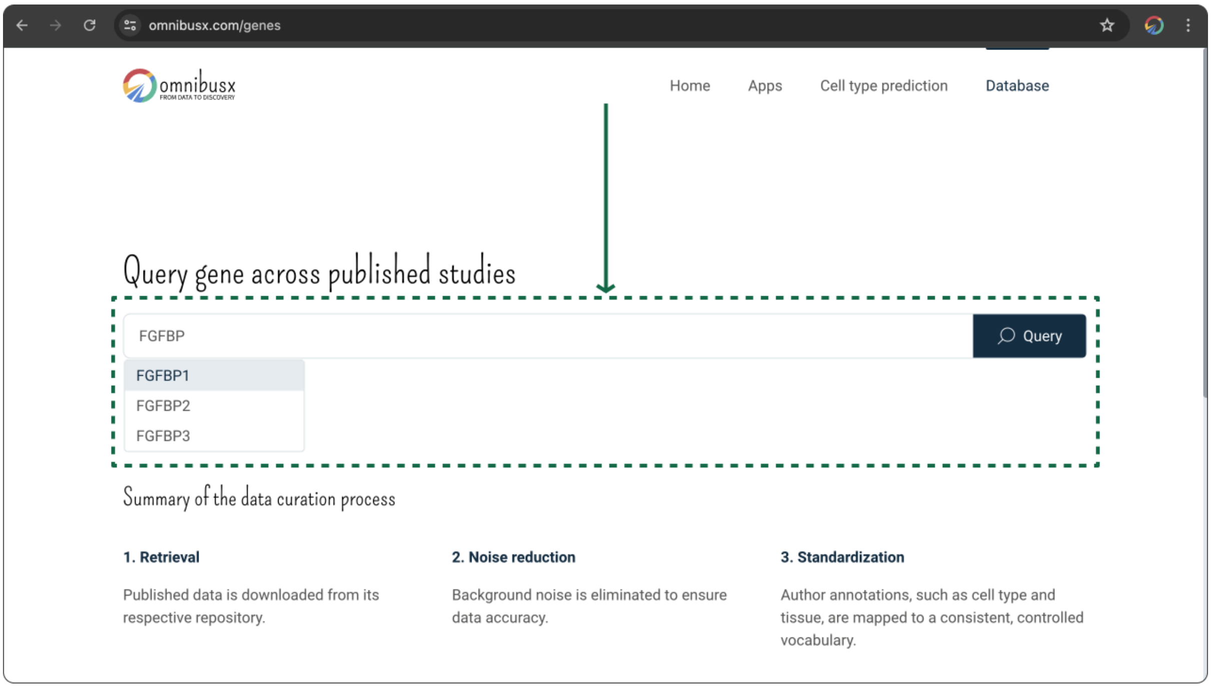


#### 2. Explore

After triggering the search, all results related to the queried gene will be retrieved and presented in two formats: table and visualization plot. In each plot, each cell population is represented by a dot, with the dot color indicating the number of supporting studies and the dot size reflecting the average expression coverage of the gene from those studies. Gene expression interpretations can be summarized as follows:

- `Specificity`: The fewer cell types or tissues a gene is expressed in, the more specific it is considered.
- `Reliability`: The greater the number of supporting studies, the more reliable the expression data.
- `Capture rate`: Higher expression coverage suggests that a gene is characteristic of certain cell types or tissues; conversely, low coverage may suggest the presence of a sub-population or limitations in detection technologies.

#### 2.1. Gene expression across cell types

All cell types and subtypes that express the queried gene are visualized using the ontology from Cell Ontology. A complex ontology structure indicates widespread gene expression across cell types, while a simple structure suggests greater specificity.

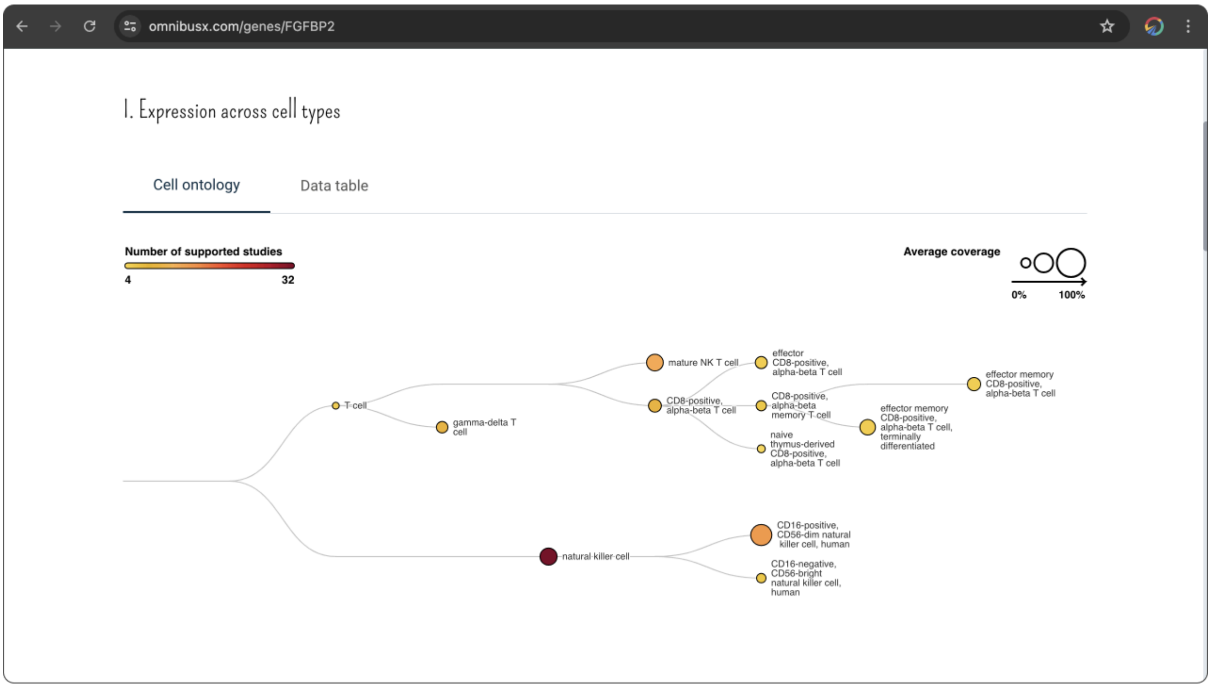


From the result table, you can access all supported studies for the expression of the queried gene in a cell type by clicking on the `Explore` button. You can also click on the cell type name to directly jump to the `Cell types search` function of OmnibusX to further study the characteristics of the cell type.

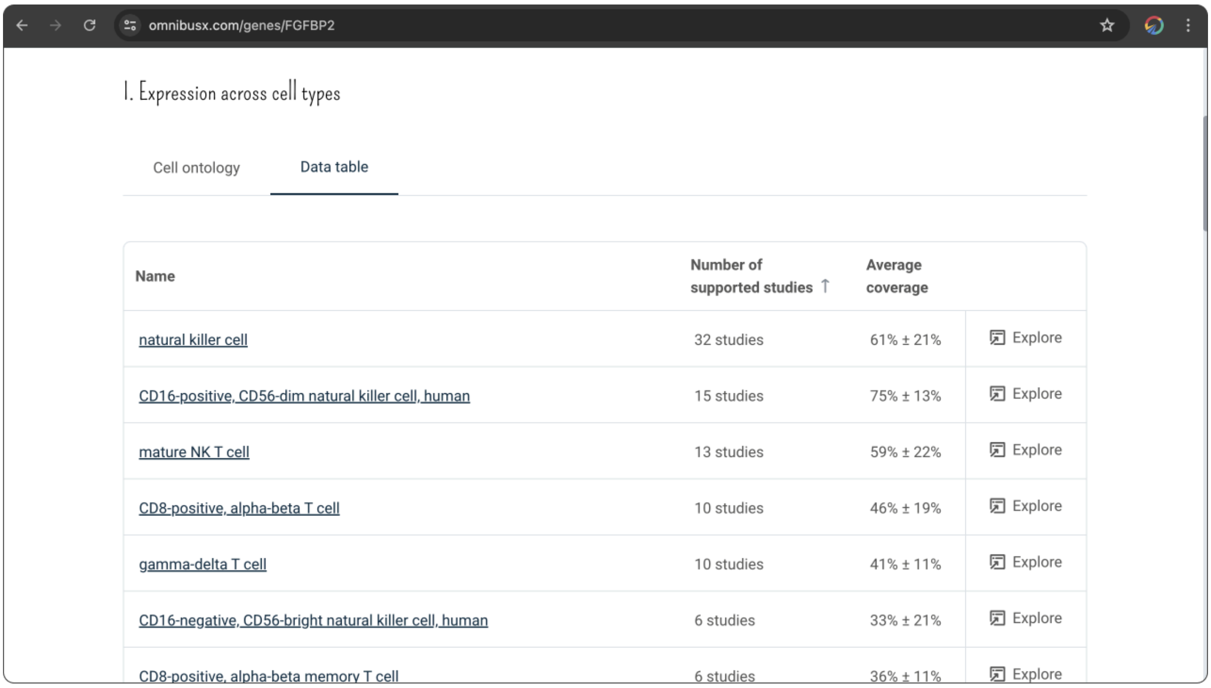


#### 2.2. Gene expression across tissues

Tissues are sorted by the number of supporting studies for the expression of the queried gene. Tissues expressing the queried gene are visualized along with all other tissues to depict the tissue specificity of the gene.

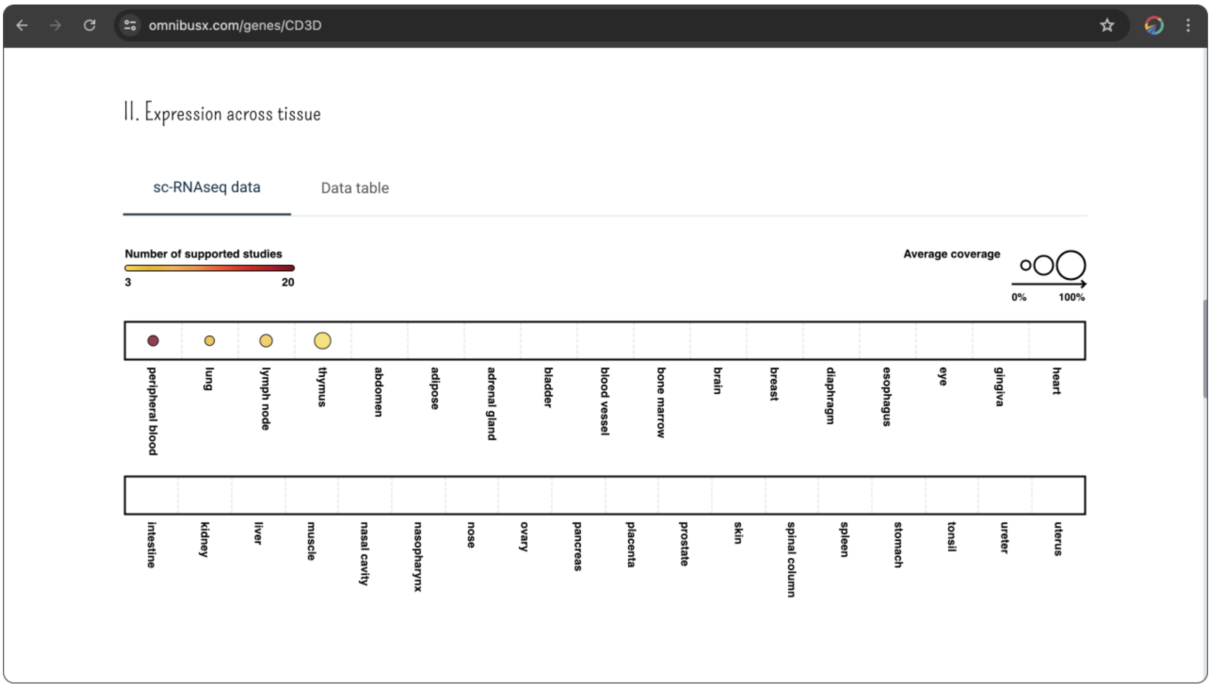

From the result table, you can access all supported studies for the expression of the queried gene in a tissue by clicking on the `Explore` button.

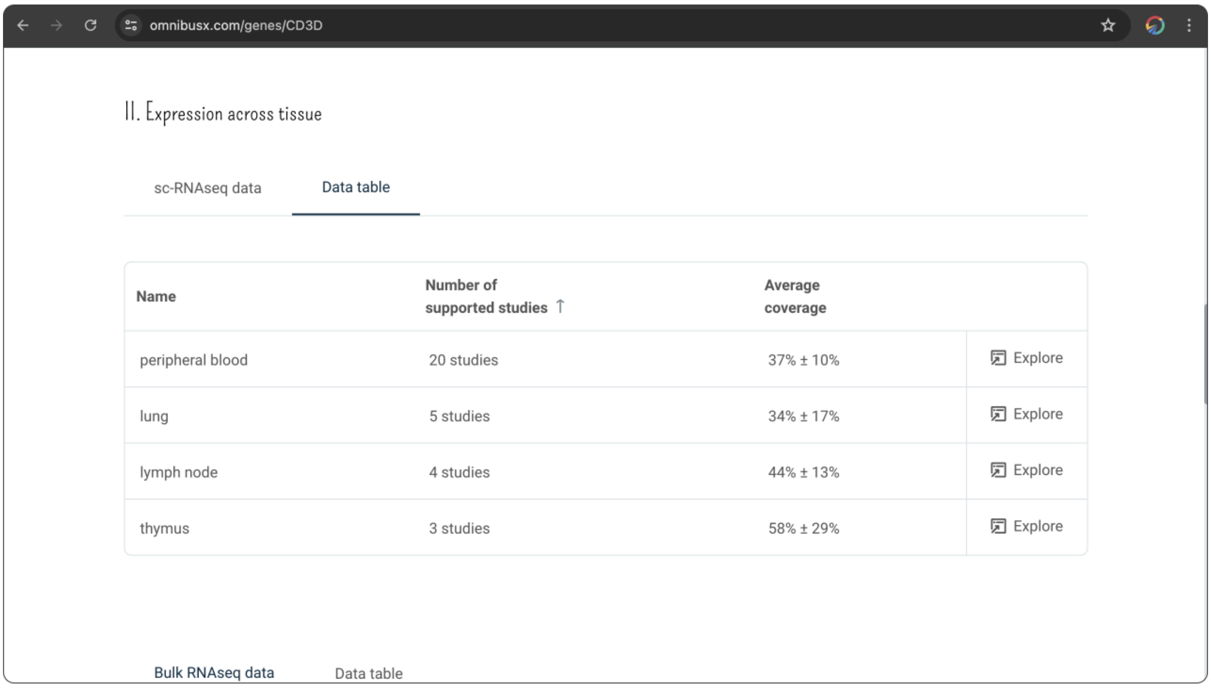


#### 2.3. Gene expression from bulk RNA-seq data

In addition to expression data from single-cell databases, we also retrieve gene expression from bulk RNA-seq datasets, including GTEx for normal tissue and TCGA for cancer tissue. This helps compare the expression of the gene between conditions and sequencing platforms to avoid biased conclusions.

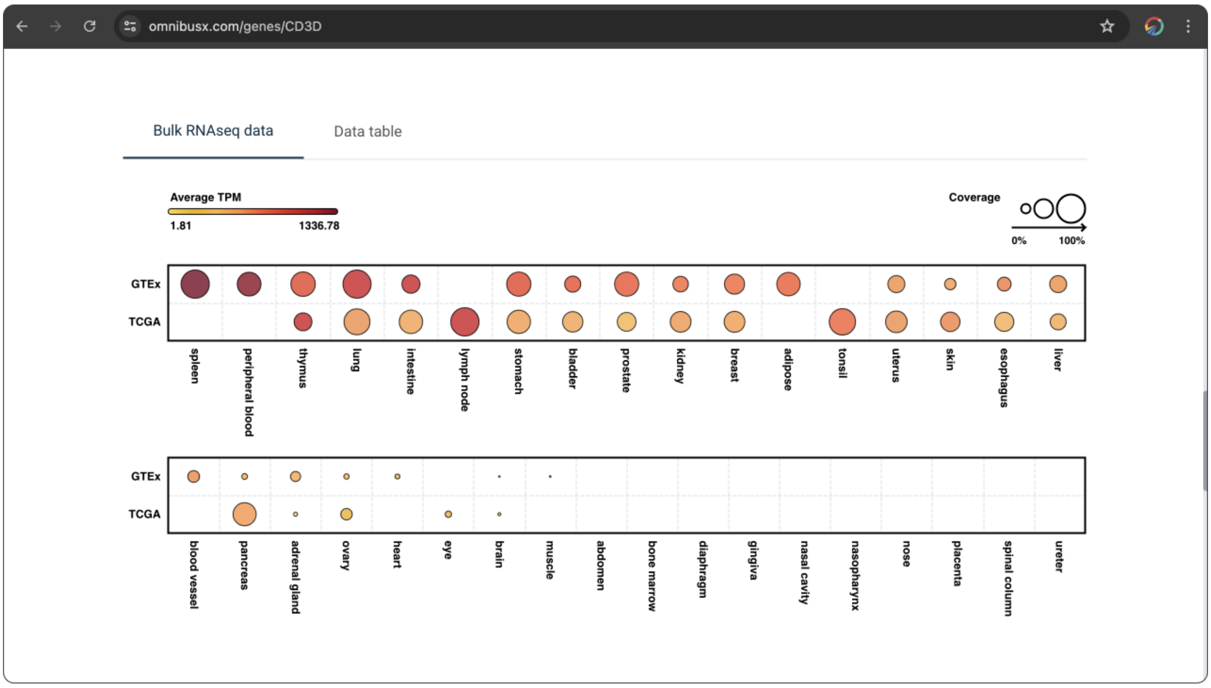


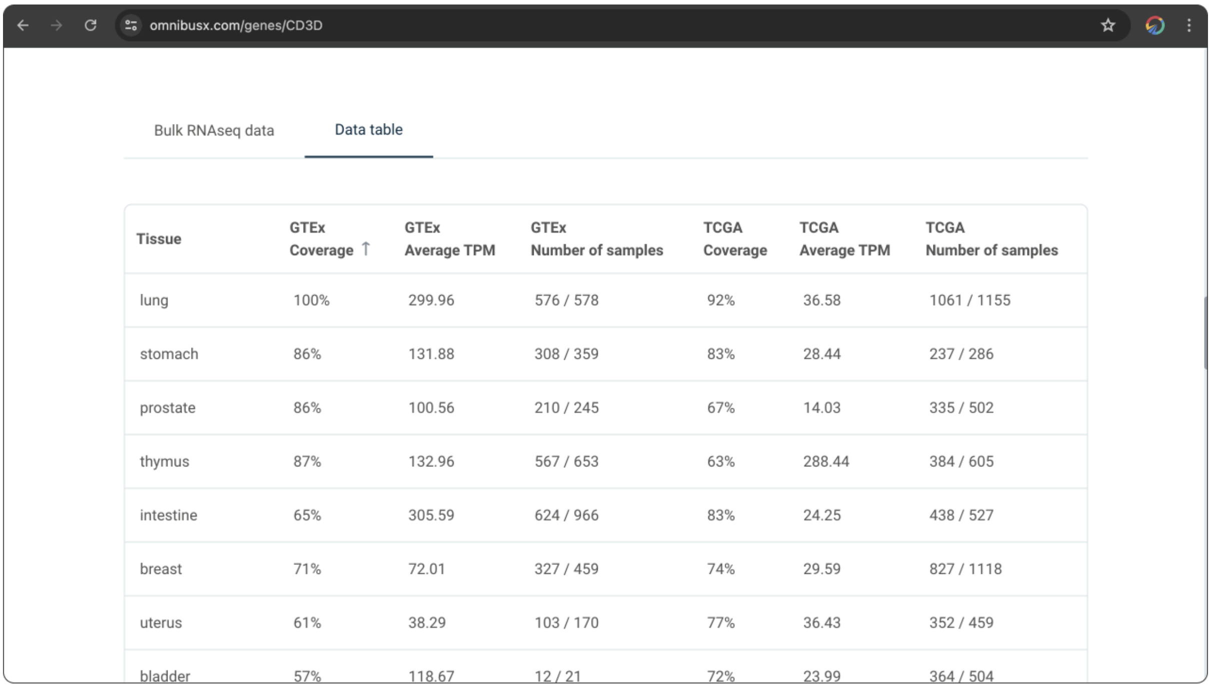


| Tissue | GTEx Coverage ↑ | GTEx Average TPM | GTEx Number of samples | TCGA Coverage | TCGA Average TPM | TCGA Number of samples |
|---|---|---|---|---|---|---|
| lung | 100% | 299.96 | 576 / 578 | 92% | 36.58 | 1061 / 1155 |
| stomach | 86% | 131.88 | 308 / 359 | 83% | 28.44 | 237 / 286 |
| prostate | 86% | 100.56 | 210 / 245 | 67% | 14.03 | 335 / 502 |
| thymus | 87% | 132.96 | 567 / 653 | 63% | 288.44 | 384 / 605 |
| intestine | 65% | 305.59 | 624 / 966 | 83% | 24.25 | 438 / 527 |
| breast | 71% | 72.01 | 327 / 459 | 74% | 29.59 | 827 / 1118 |
| uterus | 61% | 38.29 | 103 / 170 | 77% | 36.43 | 352 / 459 |
| bladder | 57% | 118.67 | 12 / 21 | 72% | 23.99 | 364 / 504 |

#### 2.4. Literature review

Other literature information about the gene, retrieved from UniProt, is also presented to support conclusions and provide explanations for observed gene expression patterns.

omnibusx.com/genes/CD3D

##### III. Literature review

[source]

| | |
|---|---|
| **Gene name** | CD3D |
| **Protein name** | T-cell surface glycoprotein CD3 delta chain (T-cell receptor T3 delta chain) (CD antigen CD3d)<br>CD3 delta<br>T-cell surface glycoprotein CD3 delta chain (T-cell receptor T3 delta chain) |
| **Synonyms** | T3D<br>hCG_40222 |
| **Description** | FUNCTION: Part of the TCR-CD3 complex present on T-lymphocyte cell surface that plays an essential role in adaptive immune response. When antigen presenting cells (APCs) activate T-cell receptor (TCR), TCR-mediated signals are transmitted across the cell membrane by the CD3 chains CD3D, CD3E, CD3G and CD3Z. All CD3 chains contain immunoreceptor tyrosine-based activation motifs (ITAMs) in their cytoplasmic domain. Upon TCR engagement, these motifs become phosphorylated by Src family protein tyrosine kinases LCK and FYN, resulting in the activation of downstream signaling pathways . In addition of this role of signal transduction in T-cell activation, CD3D plays an essential role in thymocyte differentiation. Indeed, participates in correct intracellular TCR-CD3 complex assembly and surface expression. In absence of a functional TCR-CD3 complex, thymocytes are unable to differentiate properly. Interacts with CD4 and CD8 and thus serves to establish a functional link between the TCR and coreceptors CD4 and CD8, which is needed for activation and positive selection of CD4 or CD8 T-cells. . |
| **Accessions** | B0YIY4 |

### Thank you!

We extend our heartfelt gratitude to all users of the OmnibusX Studies Search platform. Your engagement and feedback are invaluable to us and are what drive continuous improvement and innovation within our database. We are committed to supporting the scientific community by providing robust tools that facilitate groundbreaking research and discovery.

If you have suggestions, feedback, or would like to share how OmnibusX has assisted in your research endeavors, please do not hesitate to reach out to us at. Your stories inspire us, and your feedback helps us refine our tools to better serve your needs.
