## Supplementary Data 3: Cell types search tutorial for "Integrative analysis of single-cell gene expression: A comprehensive database approach"

User guide for OmnibusX web tools

### Introduction

Single-cell RNA sequencing (scRNA-Seq) technologies have revolutionized the field of single-cell biology, providing unprecedented insights into the complexities of biological systems. However, the rapid growth of data generation has presented significant challenges in data management and utilization. Researchers often face difficulties in accessing and integrating vast datasets, requiring extensive computational resources and coding effort.

To address these challenges, we have developed a comprehensive database with advanced computational tools that efficiently index massive datasets, enabling swift access to author annotations and detailed cell expression profiles. Our user-friendly interface allows researchers to quickly retrieve cell type-specific markers and characteristics, significantly reducing the time required for data processing.

The Cell Type Search tool, powered by our extensive database, helps identify cell type-specific markers and characteristics across the whole database. This guide will help you fully leverage the capabilities of the Cell Type Search tool to enhance your research outcomes.

### Cell types search

You can access the search tool directly at https://omnibusx.com/celltypes.

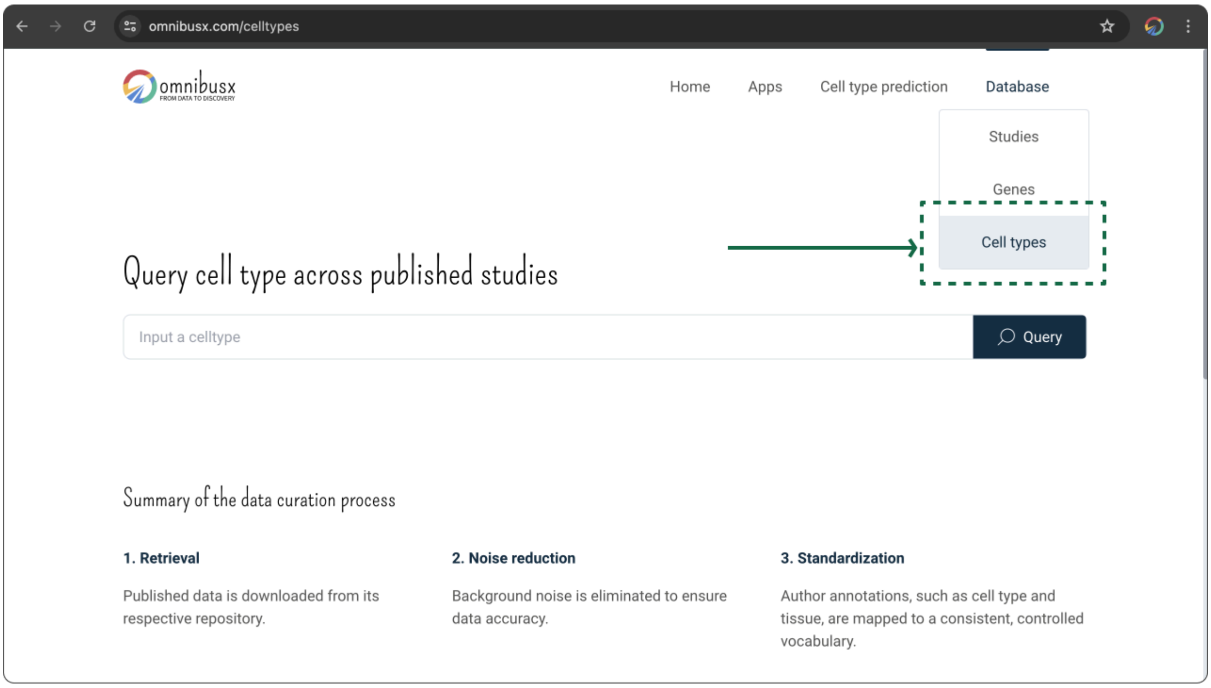


#### 1. Queries

You can begin by entering the target cell type name into the input box on the `Cell type search` page. As you type, OmnibusX will dynamically suggest related cell type names, allowing you to select the correct one directly from the dropdown menu.

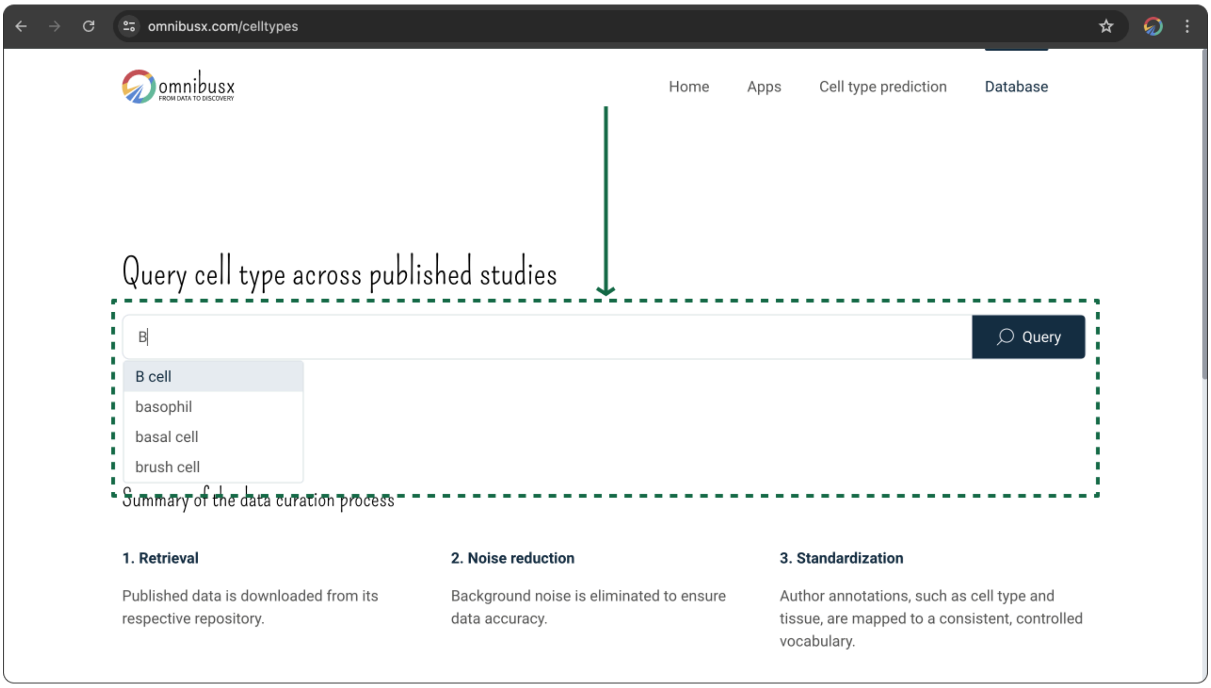


### 2. Explore

#### 2.1. Cell type lineage

All available subtypes of the queried cell types from the database are retrieved, illustrating their relationships using Cell Ontology as a sub-ontology. This visualization provides insights into the lineage and hierarchy of cell types.

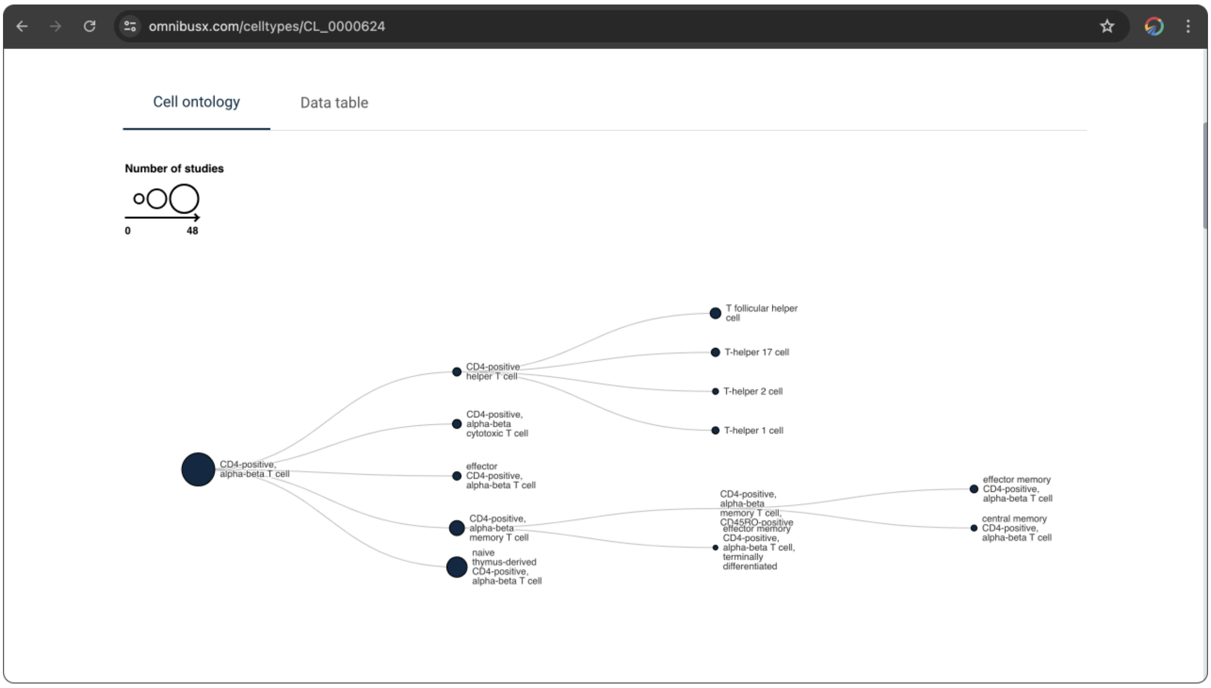


From the result table, you can access a list of studies that contain each cell subtype by clicking on the `Explore` button associated with each subtype. By clicking on a cell type ID in the table, you can trigger a detailed query for information specific to that cell type.

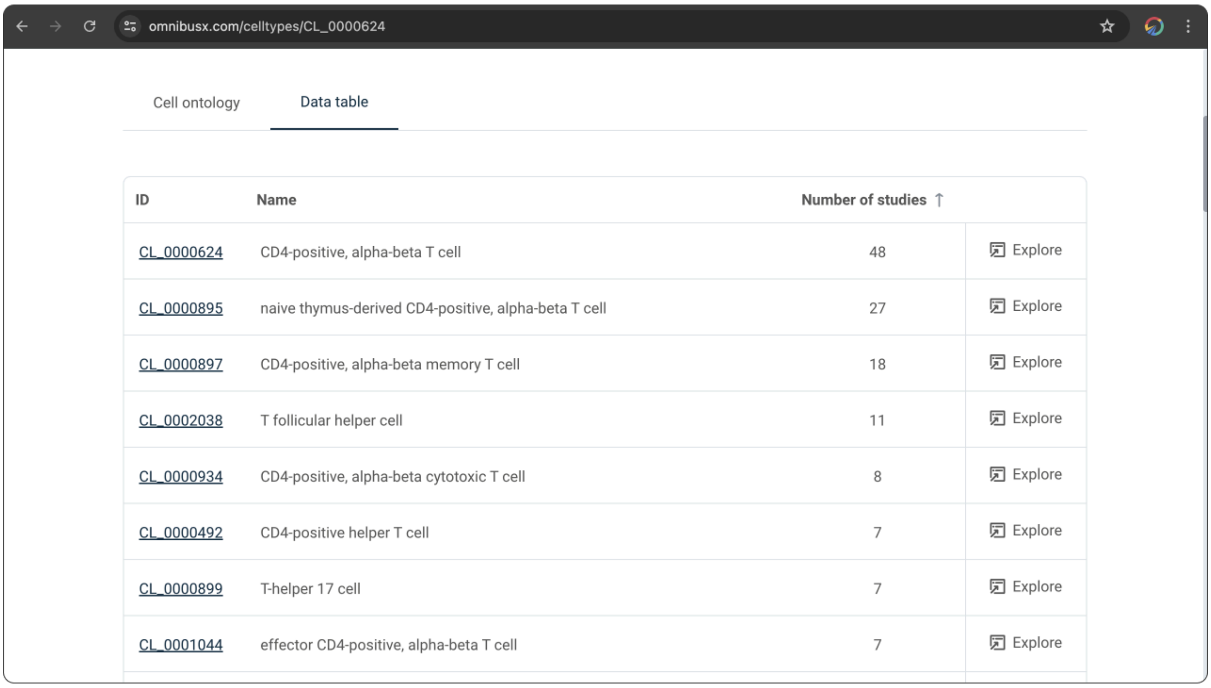


| ID | Name | Number of studies | |
|---|---|---|---|
| CL_0000624 | CD4-positive, alpha-beta T cell | 48 | Explore |
| CL_0000895 | naive thymus-derived CD4-positive, alpha-beta T cell | 27 | Explore |
| CL_0000897 | CD4-positive, alpha-beta memory T cell | 18 | Explore |
| CL_0002038 | T follicular helper cell | 11 | Explore |
| CL_0000934 | CD4-positive, alpha-beta cytotoxic T cell | 8 | Explore |
| CL_0000492 | CD4-positive helper T cell | 7 | Explore |
| CL_0000899 | T-helper 17 cell | 7 | Explore |
| CL_0001044 | effector CD4-positive, alpha-beta T cell | 7 | Explore |

#### 2.2. Cell type markers

The top 30 potential cell type markers, pre-calculated across the entire database, are displayed in a tabular format. This table includes:

- `Number of supporting studies`: Indicates how many studies have observed the cell type expresses the gene.
- `Average coverage`: Shows the average percentage of cells expressing the gene across supported studies.
- `Other major groups`: Number of other major cell types that also express the gene.
- `Other cell types`: Number of other cell types in the same major group with queried cell type that also express the gene.

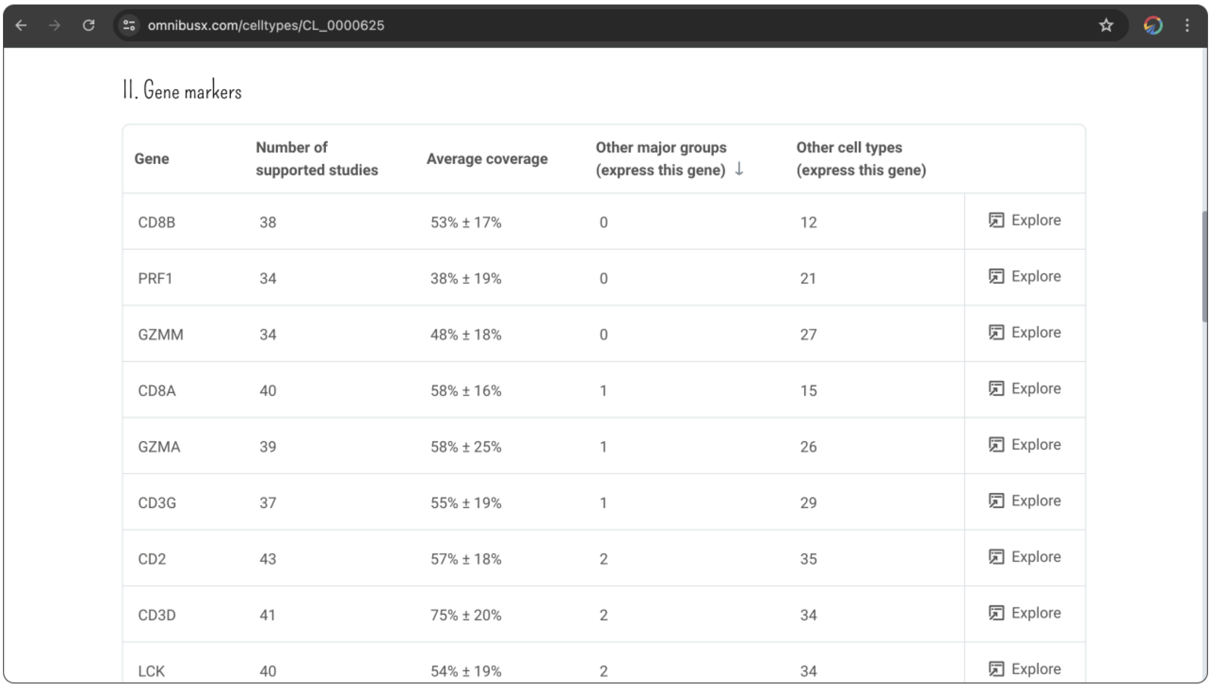


| Gene | Number of supported studies | Average coverage | Other major groups (express this gene) ↓ | Other cell types (express this gene) | |
|---|---|---|---|---|---|
| CD8B | 38 | 53% ± 17% | 0 | 12 | Explore |
| PRF1 | 34 | 38% ± 19% | 0 | 21 | Explore |
| GZMM | 34 | 48% ± 18% | 0 | 27 | Explore |
| CD8A | 40 | 58% ± 16% | 1 | 15 | Explore |
| GZMA | 39 | 58% ± 25% | 1 | 26 | Explore |
| CD3G | 37 | 55% ± 19% | 1 | 29 | Explore |
| CD2 | 43 | 57% ± 18% | 2 | 35 | Explore |
| CD3D | 41 | 75% ± 20% | 2 | 34 | Explore |
| LCK | 40 | 54% ± 19% | 2 | 34 | Explore |

A marker is considered more reliable if it is supported by a higher number of studies and has a higher average expression coverage. A marker is deemed more specific if fewer other cell types and subtypes express the gene. By clicking on the `Explore` button next to each marker, you can access the list of studies that support the expression of the marker on the queried cell type.

#### 2.3. Literature review

Other literature information about the cell type, retrieved from Cell Ontology, is also presented to provide more detailed information on the queried cell type.

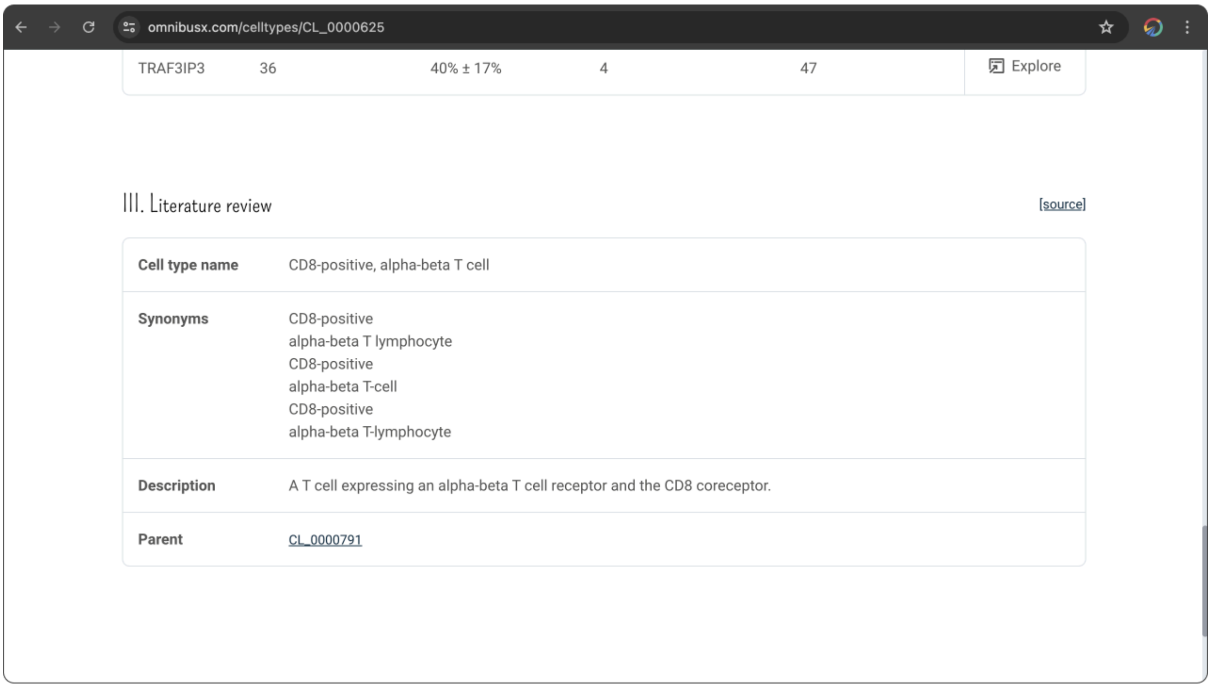

### Thank you!

We extend our heartfelt gratitude to all users of the OmnibusX Studies Search platform. Your engagement and feedback are invaluable to us and are what drive continuous improvement and innovation within our database. We are committed to supporting the scientific community by providing robust tools that facilitate groundbreaking research and discovery.

If you have suggestions, feedback, or would like to share how OmnibusX has assisted in your research endeavors, please do not hesitate to reach out to us at. Your stories inspire us, and your feedback helps us refine our tools to better serve your needs.
